# Aligning Neural Population Patterns Facilitates Motor Learning Transfer

**DOI:** 10.1101/2025.05.16.654614

**Authors:** Xiang Zhang, Zhiwei Song, Xiang Shen, Shuhang Chen, Yifan Huang, José C. Príncipe, Yiwen Wang

## Abstract

Motor learning transfer, the ability to apply skills acquired in one task to enhance performance in a related task, is driven by changes in neural ensemble activities. However, the long-term evolution of neural population dynamics during motor learning transfer remains unclear. Specifically, how do the neural patterns reorganize and stabilize over an extended learning period in the new task? To investigate the neural mechanisms in the motor cortex that enable such transfer, we employed a Brain-Machine Interface (BMI) paradigm in rats. In the experiment, rats first mastered a lever-pressing task before proceeding to a more complex but related lever-discrimination task. We analyzed neural ensemble activities by projecting them into a low-dimensional linear feature space that captures the most prominent dynamic structure. Within this space, we represented neural patterns as clusters and developed an iterative method to align similar clusters between the two tasks. Leveraging this alignment, we introduced a novel decoding approach, Clustering Alignment-based Transfer on Kernel Reinforcement Learning (CATKRL), which utilizes parameters learned from the initial task to enhance efficiency in the new task. Our results revealed that neural patterns for learned actions form distinct clusters with consistent shapes and centroid distances across tasks, and these patterns exhibit rotational evolution during learning. By integrating the cluster alignment mechanism into the RL decoder, we achieved a faster training speed with less data in the lever-discrimination task. Our results suggest that aligning neural pattern clusters can enhance BMI decoding efficiency by leveraging consistent neural representations. This approach not only provides valuable insights into the brain mechanisms underlying motor learning transfer but also holds promise for advancing multi-task learning in neuroprosthetics.

---

Motor learning ^1^ plays an important role in the development of motor behavior, as well as the acquisition of new movement skills. Through practice, individuals can adapt to novel situations and improve their performance ^2^. Motor learning transfer, the ability to apply skills learned in one task to improve performance in a related task, is a key mechanism that helps individuals increase their learning speed by leveraging previous experience ^3-6^. For example, a tennis player can learn badminton more quickly due to similar movements ^7^. Such behavioral evidence highlights the impact of motor learning transfer in accelerating skill development.

The motor cortex plays an important role in motor learning ^8^, where repeated practice can lead to specific movement representations encoded in neural firing patterns ^9^. This neural adaptability is crucial not only for acquiring new skills but also for transferring those skills to related tasks. For example, the authors ^6^ demonstrate that the covert learning in motor cortex can transfer to overt reaching behavior. Understanding how neural patterns evolve within the motor cortex is crucial to elucidating the brain mechanisms underlying motor learning.

Brain-Machine Interface (BMI) ^10-12^ technology offers a powerful platform for investigating neural patterns. By translating neural signals into action commands such as controlling a robotic arm or moving a cursor on a screen, BMI enables a concurrent recording of behavior and corresponding neural activities. This capability provides a direct observation of the brain’s internal electrical processes during task learning, allowing correlating specific behaviors learning transfer with dynamic neural patterns.

Previous BMI studies have revealed that the neural patterns during learning tend to reside within a low-dimensional subspace ^13-15^. Additionally, researchers posit that the motor learning transfer originates from a common neural substrate, which consists of motor cortical preparatory activity that facilitates the transfer of learning ^6^. However, these studies primarily focused on the neural activity changes within a single task, leaving the long-term neural population dynamics across tasks unexplored.

Analyzing neural population dynamics over extended periods for different tasks poses several significant challenges. First, the high dimensionality of the data, stemming from the large number of recorded neurons, complicates direct analysis in the original data space and brings interpretative difficulties. Second, neurons may appear or disappear across tasks due to factors such as cell death, migration, or variations in recording quality, which impedes the establishment of consistent neural correspondences, hindering cross-task comparisons. Third, the non-stationarity of neural firing patterns, due to adaptation or learning transfer, poses a significant obstacle to consistently tracking and comparing neural dynamics across different tasks. These challenges not only complicate the analysis of neural population dynamics but also impede the development of efficient decoding algorithms for BMI. To address these obstacles, we propose a novel method: Clustering Alignment-based Transfer on Kernel Reinforcement Learning (CATKRL). Our approach examines the evolution of neural patterns during motor learning transfer across related tasks and harnesses these patterns to improve decoding efficiency. The method consists of three key steps. First, we project the neural ensemble activities into a low-dimensional linear feature space that captures the most prominent dynamic structure. Then, the neural patterns can be represented by neural pattern clusters rather than individual neurons. Third, we propose to iteratively align similar neural clusters between two tasks in this linear space. In this way, the learned neural-action mapping in the initial task could be reused in the related new task, which could facilitate the learning efficiency in the decoding process.

We evaluate our proposed algorithm through an experimental paradigm where rats first master a lever-pressing task before progressing to a more complex lever-discrimination task. The second task builds upon the motor skills developed in the initial task. We expect that the rats can leverage their prior experience from the lever-pressing task to enhance the speed of learning in the lever-discrimination task. We simultaneously recorded the behavior and neural ensemble activities throughout the learning process of both tasks, aiming to uncover the dynamic evolution of neural patterns along with behavior improvement. Furthermore, we examine whether our proposed algorithm can effectively detect and harness these evolving neural patterns to facilitate motor learning transfer. Specifically, we seek to determine if our method enables faster learning of the second task compared to learning from scratch and whether it can achieve comparable performance with reduced training data.

## Results

### Behavioral Transfer across Tasks

Learning a complex task can be decomposed into a sequence of simple tasks ^16,17^, such that subjects are able to manage and re-utilize the experience from the simpler task to speed up learning. We investigated this motor learning transfer mechanism in BMI experiment involving SD rats. The rats were trained to perform two related tasks over days of training. Task 1 was relatively easy as shown in Fig. 1a, the rat stayed inside a behavior box and waited for an audio cue as a trial start. It was required to press and hold the lever for 0.5s to get water reward. After the rats were well-trained in task 1, they were instructed to do task 2 with more complex movements as shown in Fig. 1b. The rats needed to differentiate between two audio frequencies in each trial and hold the lever corresponding to the specific frequency. The experience of pressing the same lever in the first task is expected to be reutilized by the rats to achieve motor learning transfer.

**Fig. 1.**
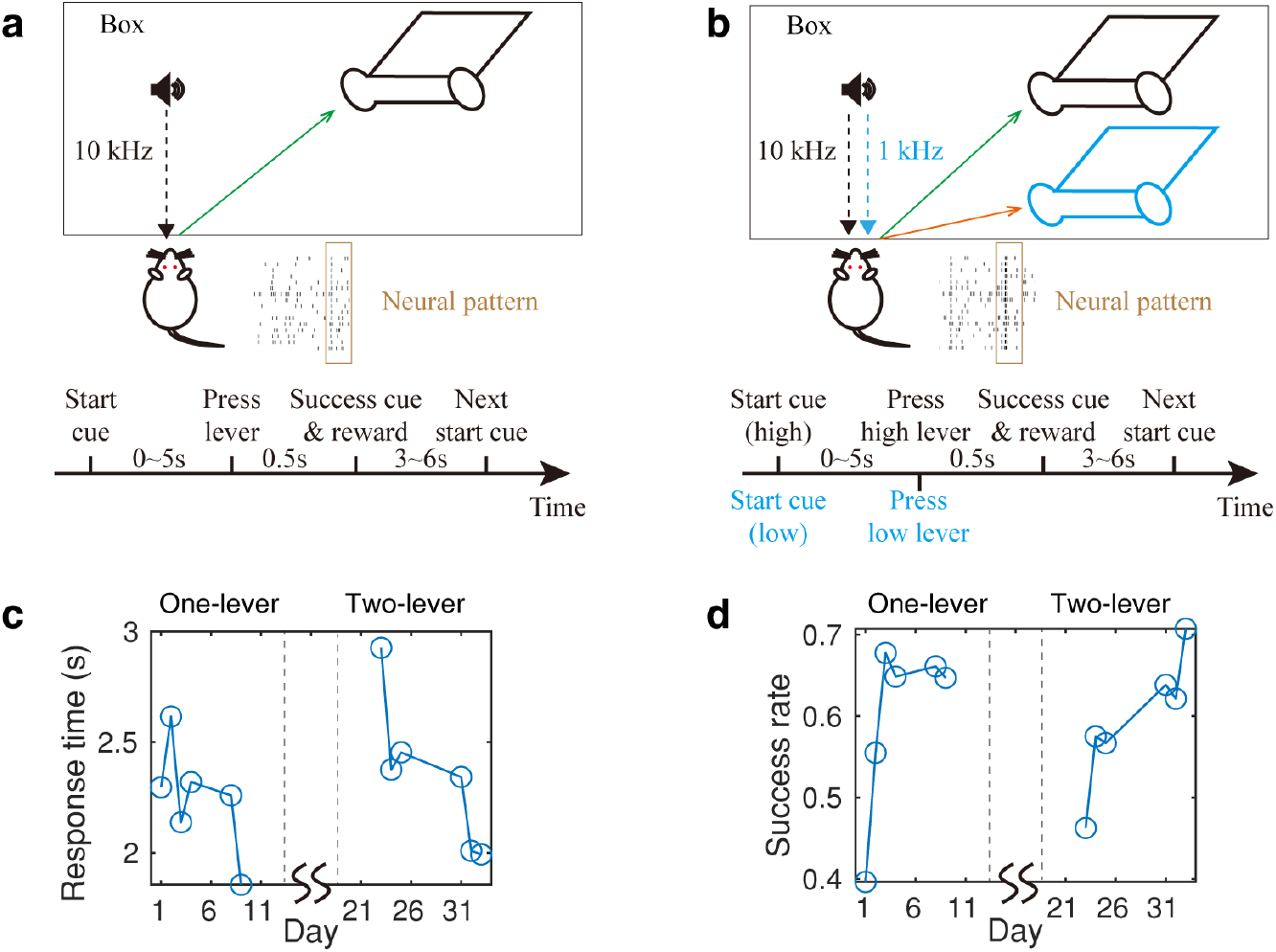
Experimental paradigms that enable the motor learning transfer of SD rats. **(a)** The lever-pressing task for SD rats. When the rat heard an audio cue (10 kHz) from the speaker, it should press and hold the lever for 0.5 s within 5s to get the water reward. The timeline shows the task flow. **(b)** The two-lever discrimination task. The rat was trained on the one-lever task in the first half of the experiment. After the rat was proficient in the one-lever task, it was trained on the two-lever discriminative task. The second task for rats is related but more complex than the first task in **a.** The rat needed to distinguish two sound frequencies and choose the correct lever to get the water reward. The timeline shows the task flow. **(c)** The average response time of one rat over the training period. The horizontal line represents the days that the rat was trained. The vertical line shows the average response time across trials on each day. The response time is calculated as the interval between the trial start cue and the successful cue. **(d)** The success rate of the rat’s task over the training period: the number of successful trials divided by the total number of trials on each day. The other conditions are the same as **c**.

We first measured the rat’s performance of behavioral learning in the one-lever pressing and the two-lever discriminative task. The goal is to verify that, at the behavioral level, the motor learning transfer occurred during the experiment. We focused on the response time and the success rate of the rats to measure the transfer effect. After 10 days training, the rat was proficient at the one-lever task, as shown by the low response time (∼2 s) and the high success rate (∼70%) in Fig. 1c and 1d respectively. Then the rat started the training to learn the two-lever discriminative task. In the beginning, the rat was not very familiar with the new task, which is shown by the high response time (∼3 s) and the low success rate (∼50%). After around 10 days of learning, the rat could reach a similar task performance (∼2 s response time and ∼70% success rate) as the one-lever pressing task as shown in Fig. 1c and 1d. Even though task 2 is more complex compared with task 1, the rat could master the second task using a similar amount of time (∼10 days) as the first task, which demonstrates that the experience in the first task helped the rat to learn the second task. This suggests that the motor learning transfer occurred on the rat at the behavioral level in this BMI experimental paradigm.

The learning transfer at the behavioral level implies that there should be commonalities at the neural level inside the brain. Existing literature also posits that a certain static neural feature space persists invariantly during learning ^13,18^. While numerous mechanisms may underpin the connection between simple and complex tasks in the brain, our hypothesis is relatively simple: despite potential variability in individual neuron activities across tasks, a stable and distinct representation of learned actions is preserved within the neural ensemble space.

To further investigate the hypothetical strategy of learning transfer, a quantitative analysis of action representation at both individual and ensemble neural levels is required. At the single neuron level, we treat neural tuning as a statistical representation of individual neural activity. The change of tuning means the alteration in the average firing rate associated with each action. Previous studies suggest that the neural tuning preferences for different actions change over time ^19-22^. In line with these findings, we also observed neural tuning changes across tasks in our experiment.

At the neural ensemble level, we consider Fisher information ^23^ as the metric for action discrimination ability. Fisher information is inversely proportional to the minimal discernible action change ^24^. Consequently, it can serve as an indicator of the capacity to distinguish different actions from the neural ensemble response. Within a local space of neural ensemble activities, the neural patterns for the same action form a cluster. In this case, the Fisher information ^25^ can be related to the feature of each cluster and differences among actions. In our experiment, we identify a linear feature space to represent neural patterns, where the cluster shape of the same action and inter-cluster distance among clusters remain similar across tasks. In this feature space, the action discrimination ability is preserved during task learning. In the new task, although individual neural activity changes, the preserved discrimination capability ensures a way to align the neural patterns, establishing the basis of the learning transfer strategy.

### Simulated scenario: single neural tuning changes lead to neural ensemble pattern rotation in feature space

To further elucidate the learning transfer strategy from task 1 to task 2, a critical process is to understand the preserved discrimination ability by establishing the connection between neural ensemble and level the single neuron level. We propose to utilize the linear Fisher information ^24,25^ to characterize this relationship during task learning. Linear Fisher information represents the information that can be extracted by a locally optimal linear decoder. The discrimination capability *d*_*a*_ of the action *a* could be calculated as follows:

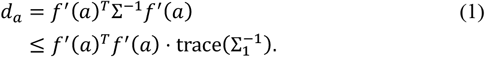

The column vector *f*(*a*) = {*f*_1_(*a*), *f*_2_(*a*), … , *f*_*N*_(*a*)} contains the normalized average firing of all *N* neurons to action *a* in task 1. 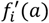is the gradient of *f*_*i*_ (*a*) with respect to action *a*. Σ_1_ is the covariance matrix of neural firing. *d*_*a*_ is upper bounded by two components: the inter-cluster distance *f*^′^(*a*)^*T*^*f*^′^(*a*) and the cluster shape approximated by Gausiasn distrubiton with variance Σ_1_. When learning task 2, if both the inter-cluster distance and the cluster shape of learneed actions remain similar, the discrimination ability will be constricted within the same upper limit. In this case, we could align the neural patterns from task 2 to task 1. This transfer learning strategy could be formulated as the following optimization problem:

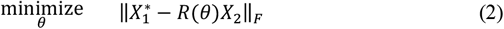

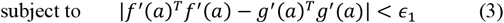

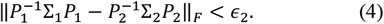

*θ* is the alignment angle between neural patterns of task 2 and 1. *R*(*θ*) ∈ ℝ^*D*×*D*^ is the rotation matrix composed from *θ. D* is the number of dimension of the feature space. 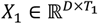 and 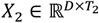 are the neural patterns in task 1 and 2 respectively. *T*_1_ and *T*_2_ is the total number of neural patterns in task 1 and 2 respectively. *X*^∗^ = {*x*|*x* ∈ *X*_1_, *x*_2_ ∈ *X*_2_, ∀*x*^′^ ∈ *X*_1_, ‖*x* − *R*(*θ*)*x*_2_‖ ≤ ‖*x*′ − *R*(*θ*)*x*_2_‖} denotes the closest neural patterns in task 1 after the alignment of task 2 patterns. ‖⋅‖_*F*_ is the Frobenius norm. The optimization goal is to align the neural patterns *X*_2_ in task 2 as close as possible to its closest patterns in task 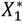.

Equation (3) and (4) are the constraints that ensure the neural ensemble activities can discriminate a specific action, establishing the prerequisites for alignment. The vector *g*(*a*) = {*g*_1_(*a*), *g*_2_(*a*), … , *g*_*N*_(*a*)} is the normalized average neural firings of all neurons to action *a* in task 2. 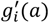 is the gradient of *g*_*i*_ (*a*) with respect to action *a. ϵ*_1_ and *ϵ*_2_ are two small threshold values. Σ_1_ and Σ_2_ are the covariance matrix obtained from *X*_1_ and *X*_2_ respectively. *P*_1_ and *P*_2_ are the eigenvector matrices of Σ_1_ and Σ_2_ respectively. Constraint (4) ensures that the inter-cluster distances are maintained, preserving the distinctiveness of learned actions. Constraint (5) means that the cluster shapes remain largely unchanged, indicating a consistent representation of the same action at neural ensemble level. When conditions (4) and (5) are satisfied, an individual neural tuning change from *f*_*i*_ (*a*) to *g*_*i*_ (*a*) might result in a rotation of the neural populational patterns. To utilize the existing neural patterns for the efficient task learning, the alignment angle *θ* from task 2 to task 1 could be calculated as follows. A more detailed iterative alignment algorithm is presented in Methods section.

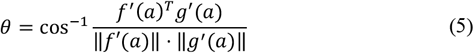

An intuitive illustration for the proposed hypothesis is presented in Fig. 2, which illustrates the hypothetical progression of neural patterns across two tasks. Tasks 1 and 2 are the simulations that imitate the one-lever and two-lever tasks respectively. In tasks 1 and 2, the Rest and Press high lever actions are shared. One of the neurons (neuron 1) changes its preferred action, while other tuning stays the same (Fig. 2a), which is a common scenario that happens during BMI experiment ^20^. Transitioning from task 1 to task 2, as shown in Fig. 2b, the centroids of the neural activities for each action change due to the tuning difference (*f*_1_(*a*) → *g*_1_(*a*)). At the same time, the inter-cluster distance (constraint (4)) and the cluster shapes (constraint (5)) and learned actions remain consistent across task, mirroring our experimental observations. When neuron 1 starts tuning more to the high press action, it results in higher firing rates of press-related cluster (green). This contributes to a rotation *θ* in Fig. 2c of the major principal direction of the learned actions connecting both rest-related cluster (red) and press-related cluster (green) in the linear feature space (grey plane). Consequently, a rotation of the neural clusters is observable in the feature space.

**Fig. 2.**
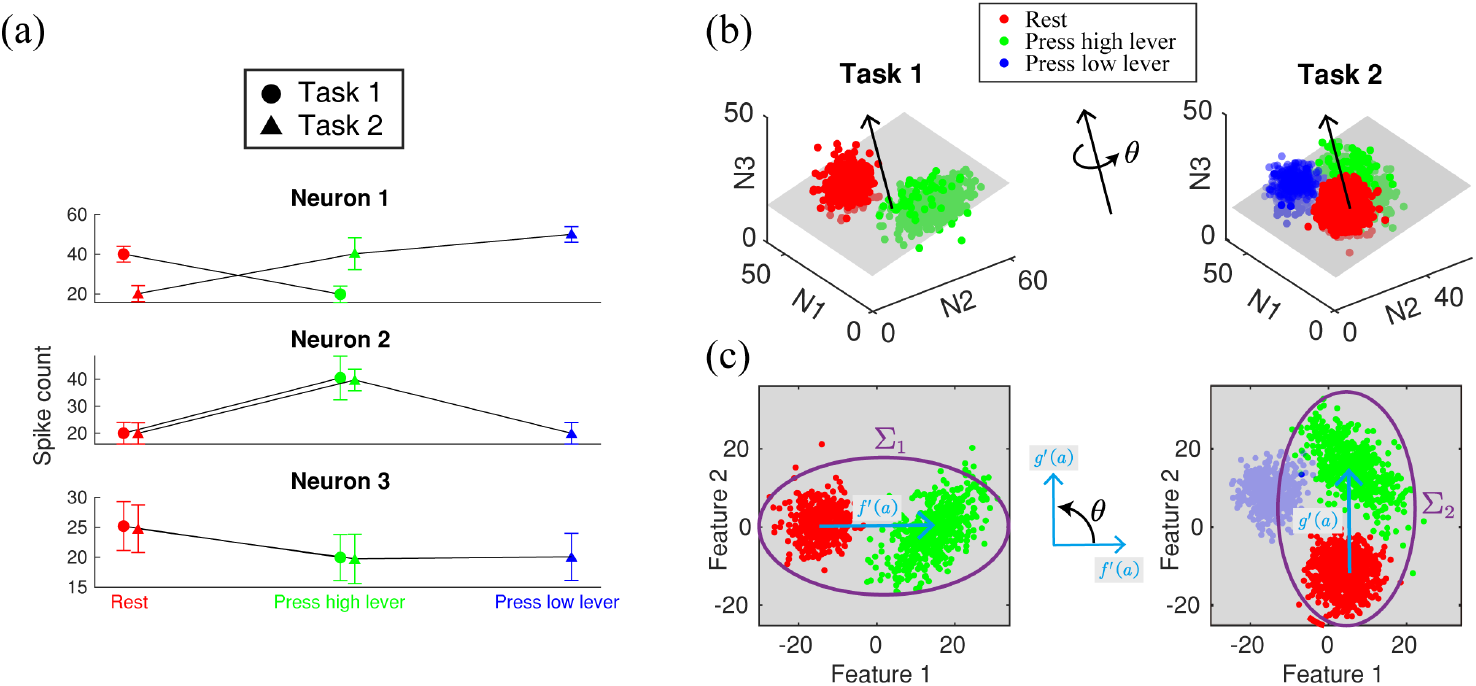
Simulated illustration of the neural pattern evolution during the learning of two tasks. **(a)** The tuning curves of simulated neurons across tasks. The horizontal axis represents two shared actions across tasks. The vertical axis represents the spike counts of each neuron at each action in the two tasks. **(b)** The spike firing patterns in the neural ensemble space of the two tasks respectively. Each axis represents the spike count of each neuron respectively. **(c)** The neural patterns linearly projected on the feature space.

The simulation presents a simplified case in which only one neuron alters its tuning can lead to the rotation of the populational patterns, while the discrimination ability of the learned actions remains the same. In real-world situations, however, the preferred movements of multiple individual neurons may shift, resulting in a more complex change in the coordinates of learned neural patterns. Nonetheless, if we can identify a linear feature space that reveals an alignment in the neural pattern, it also indicates the rotational dynamics within the original neural ensemble activity.

### Neural pattern development and algorithmic alignment iterations during motor learning transfer

To further investigate the alignment mechanism, we visualized the neural activity evolution during the real motor learning transfer experiment (Fig. 3). Since the neural activities are high-dimensional, we were motivated to examine the neural patterns in the feature space after a linear dimension reduction using jPCA ^26^ (Methods). One important property of jPCA is its ability to show the most significant rotational structure of the neural population dynamics, as shown in Fig. 3.

**Fig. 3.**
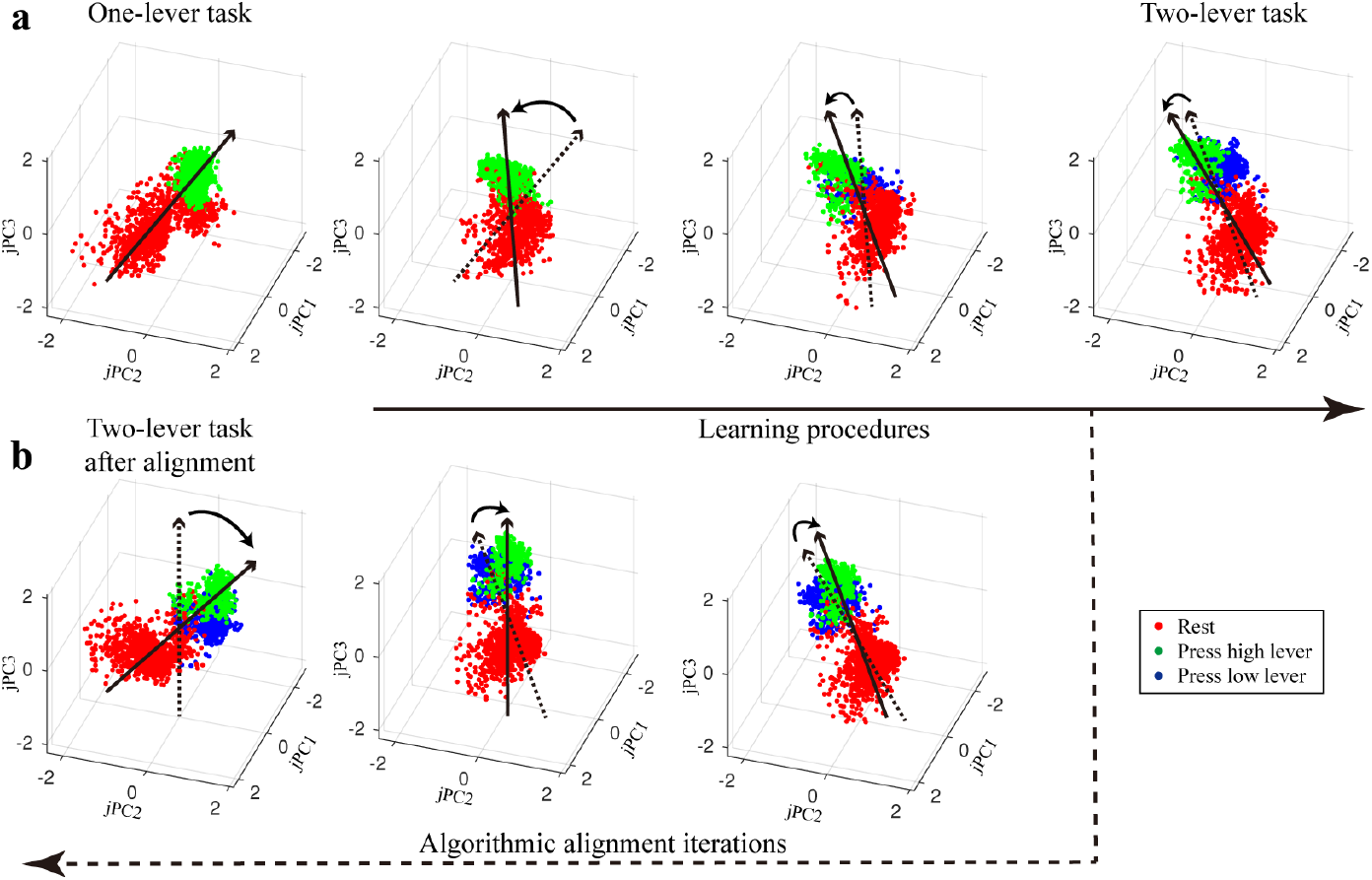
The clustering alignment method reveals the neural pattern development in the feature space. **(a)** The neural patterns in the feature space after jPCA. The dots denote neural data points in feature space and the colors correspond to different actions. The solid straight arrow indicates the direction from the centroid of the red cluster to the centroid of the green cluster. The dotted straight arrow illustrates the direction of the preceding plot, while the curved arrow portrays the rotation process. The neural pattern evolvement is shown from left to right. For each subplot, the axes represent the first three jPCA components. The red, green, and blue labels the action Rest, Press high lever, and Press low lever respectively. **(b)** The results of the clustering alignment iterations. The iteration procedure is shown from right to left. The figure conventions are the same as in **a**. The relative positions of clusters in the alignment iteration match the development of neural patterns (Methods).

One crucial observation in the low-dimensional feature space is the distinct positioning of the neural patterns of two tasks (Fig 3a, first and last subplot), and the tendency for neural data points corresponding to the same action to cluster. During the learning procedure (Fig 3a), the positions of the cluster have gradually shifted, but the shape of the clusters (red and green) remain similar, while the centroid distance also persists. Moreover, the relative position of the two clusters (solid black arrow in Fig. 3a) undergoes counterclockwise rotation. This phenomenon demonstrates the discrimination ability of the learned actions, which might allow the brain to align the current neural patterns to previously encountered patterns. This alignment could facilitate the reutilization of the learned neural-muscular mapping, which accelerates the learning of new tasks. To further quantify and utilize the alignment mechanism, we propose a computational tool to mimic the transfer procedure, exploring how the neural pattern clusters are aligned over time.

Here we propose an iterative alignment algorithm (Methods): we first cluster the neural patterns in two tasks respectively. Then we identify the neural cluster pair with the most similar shape across two tasks. Finally, we iteratively align the whole neural patterns in the feature space based on the selected cluster pair. We present the algorithmic alignment iterations from two-lever task to one-lever task in Fig. 3b. We note that the relative position of the two clusters (black arrow in Fig. 3b) rotates clockwise in the feature space, reflecting the actual neural pattern development observed in the above subplot in Fig. 3a.

### Cluster Alignment Enhances Decoding Abilities for Brain-Machine Interface

The proposed neural mechanism for motor learning transfer might also be applied to the decoding task in BMI. This could be achieved by constructing the reservoir of the learned movements in the decoder, and subsequently aligning the similar clusters. This approach could potentially endow several advantages. First, the decoder could re-utilize the learned neural-muscular mapping (parameters in the decoder) to increase the learning speed of a new but related task (learning efficiency). Second, since the decoder’s parameters are reused, the decoder would need fewer training samples in the new task (data efficiency). Lastly, the decoder could maintain its performance when switching back to the old task (memory) after undertaking the new task.

In this study, we propose a novel approach Clustering Alignment based Transfer on Kernel Reinforcement Learning (CATKRL) to enrich the learning ability of BMI. CATKRL first clusters the neural patterns within the feature space. Given the previous discovery of the rotational structure in jPCA, we also use t-SNE ^27^ to provide a non-linear feature space to better visualize the neural-action mapping. The advantage of using t-SNE lies in its ability to preverve the distribtuion of the neural activity within the feature space. The relative positions of neural activities are maintained, enabling the alignment to operate effectively. Within this feature space, CATKRL detects and aligns the most similar clusters, transfers the neural-action mapping from the existing cluster, and finally explores possible actions from the aligned neural patterns through trial and error (Methods). CATKRL was tested on the rat experiments from the simple task to the complex task.

To verify the first advantage of learning efficiency, we trained all the RL decoders from the simple one-lever pressing task to the complex two-lever discrimination task. All the decoders, a baseline RL decoder with a neural network structure (blue), the clustering KRL with no alignment (green), and CATKRL (red) have similar performance in the first task since there are no pre-trained parameters (Fig. 4a). However, when facing the second task, all three decoders’ performances drop due to the neural pattern changes. Yet CATKRL has the highest success rate during the initial stage and converges to the best performance with the least training samples. A similar phenomenon is also observed in the testing performance over multiple subjects (t-test, p<0.05) as shown in Fig. 5a.

**Fig. 4.**
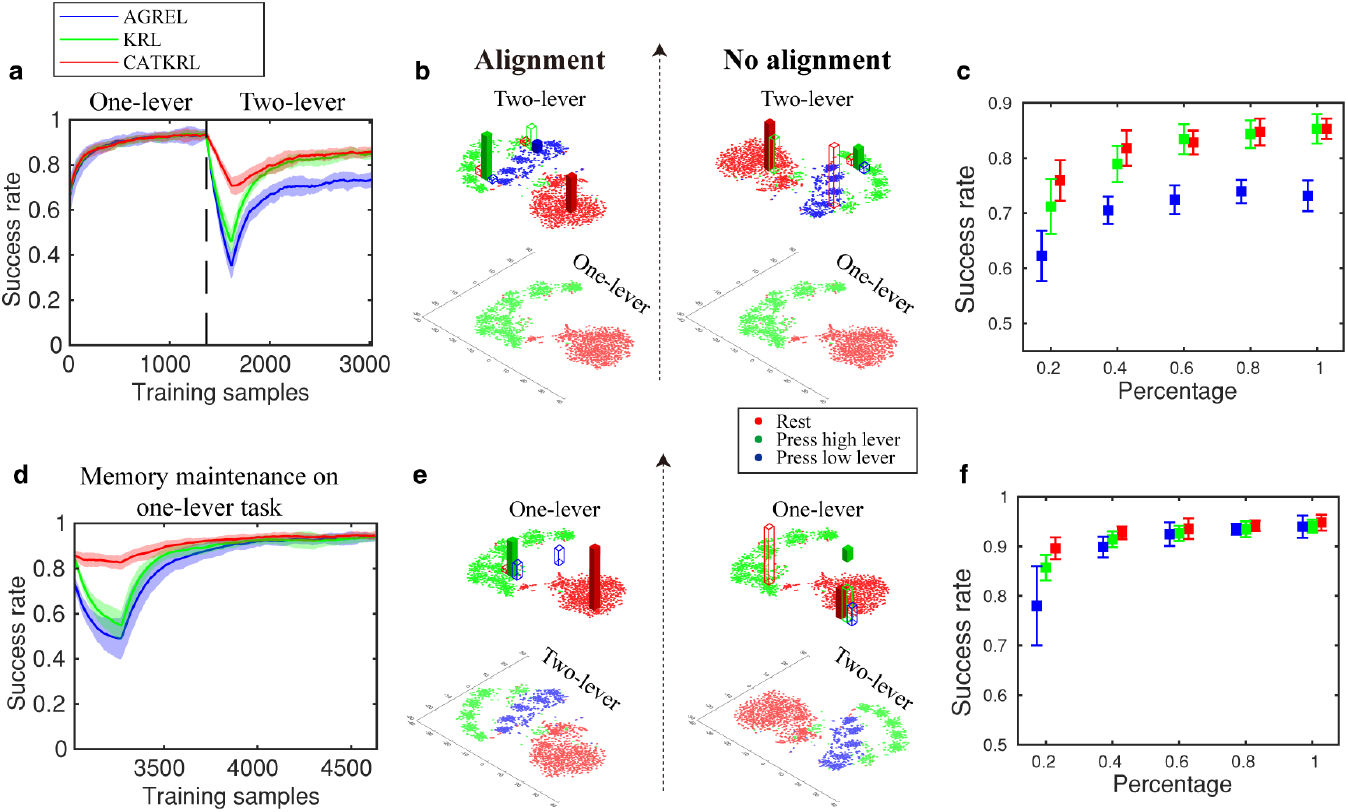
Computational validation for the motor learning transfer mechanism in BMI decoder. **(a)** The learning curve of the decoder from one-lever task to two-lever task. The x-axis is the training samples for the decoder. The y-axis is the success rate within a sliding time window (200 samples). Three decoding models AGREL (blue), Clustering KRL (green), and CATKRL (red) are tested on the same neural data. Each model’s performance is averaged over 20 data reshuffles. The solid lines show the average results, and the shaded areas represent the standard deviations. **(b)** The bottom figure represents the neural patterns in the feature space of the one-lever task, which are used to build the initial decoding parameters in the two-lever task. The color dots represent the ground truth of the action labels. The top two figures represent the neural patterns of the two-lever-discrimination task. The bars represent the action selection by CATKRL (left) and Clustering KRL with no alignment (right). The color of the bar represents a specific action chosen. The center of the bar represents the cluster centroid generated by the algorithms. The solid bar means that the algorithm has chosen the correct action of the neural pattern, which increases the decoder’s performance. In contrast, a hollow bar represents the choice of conflict actions, which decreases the decoder’s performance. The length of the bar represents the frequency of choosing the corresponding action inside each cluster during the first 50 samples. **(c)** The testing performance in the two-lever task when the decoder uses different portions of training data. The initial condition is the training convergence in the one-lever task. The x-axis represents the percentage of data in the two-lever task to be used. A smaller percentage means less training data are used for the decoder. The y-axis represents the success rate of different decoding models tested over 20 data reshuffles. The dots and the whiskers represent the mean and the standard deviation of the success rate, respectively. **(d-f)** The figures demonstrate the memory scenario transitioning from two-lever task back to one-lever task. The notations are consistent with those outlined in **a-c**.

**Fig. 5.**
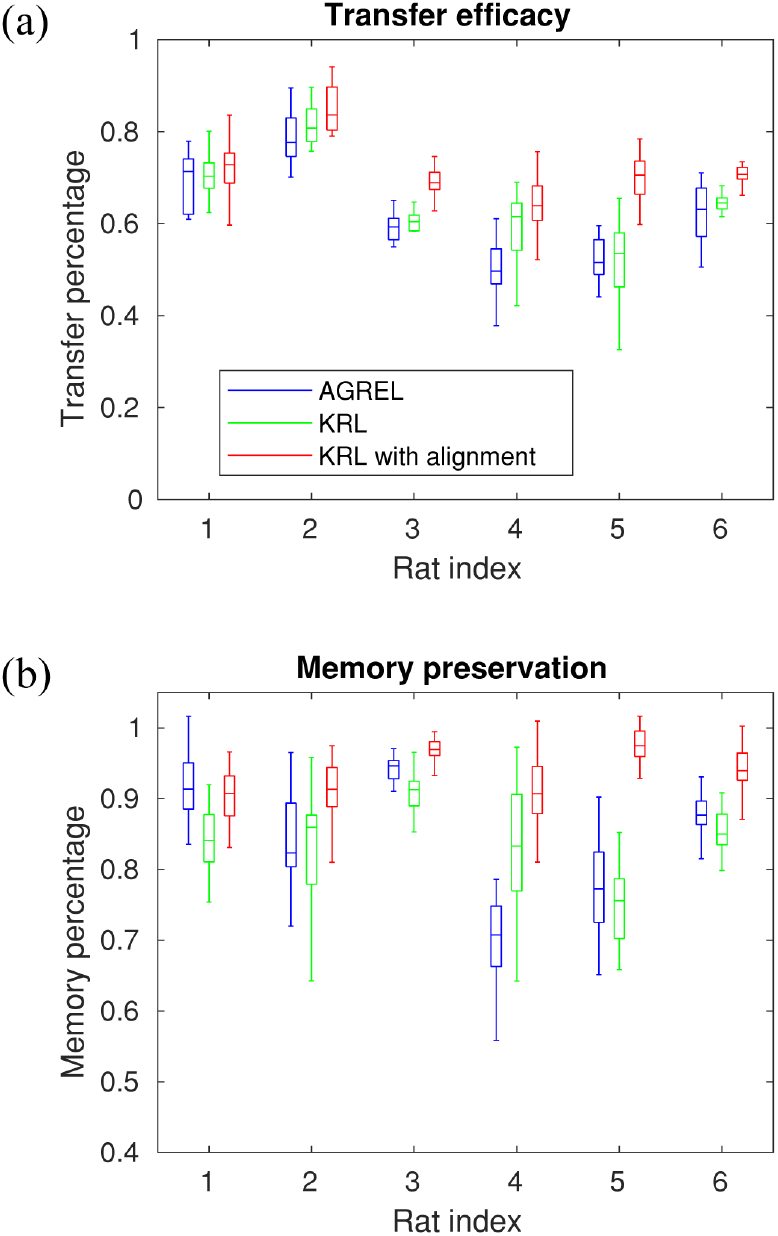
Performance across multiple subjects. **(a)** The transfer efficacy across 6 rats, computed as the success rate ratio between the early two-lever task and the convergent one-lever task. **(b)** The memory preservation across the same 6 rats, computed as the success rate ratio between the early back to one-lever task and the convergent one-lever task.

To demonstrate the reasons for the better learning efficiency of CATKRL, we present the neural data distribution and the corresponding action assignment by different decoders in Fig. 4b. Following the training in the one-lever task, both CATKRL and clustering KRL have recorded the neural patterns and the weights to the corresponding actions (as shown by the bars in Fig. 4b). When facing the two-lever task, the decoders would choose actions based on the experience obtained from the one-lever task. The absence of neural pattern alignment (right figure, Fig. 4b) led to a mismatch of the neural patterns in the two-lever task with those from the one-lever task. This discrepancy would result in a different action selection from the ground truth action labels (green hollow bars on red cluster), thereby reducing the decoding performance. As our proposed method has aligned the neural patterns in task 2 to the most similar cluster in task 1, the decoder has a high chance of choosing the correct action by utilizing the transferred neural-action mapping across two tasks (solid bar centered at neural data cloud with the same color).

This gives CATKRL a good initial performance when facing the two-lever task. These comparisons suggest that CATKRL efficiently transfers the learned neural action mapping in task 1 to increase the learning speed of the related task 2.

To verify the second advantage of data efficiency, all RL decoders were trained to have a good performance at task 1, serving as the initial condition of the second task. Then the decoding performance was tested with different amounts of training samples in task 2 (Fig. 4c). Compared with the baseline RL method (blue), CATKRL (red) consistently outperforms under different data percentages. Compared with the same method with no alignment (green), we find that CATKRL has a higher testing success rate when there are small amounts of training samples (≤ 40%). Even though the final performance values of the two algorithms are similar, CATKRL requires fewer training samples (60%) to reach peak performance. These results also suggest that CATKRL could actively re-utilize the neural patterns of the same action from the learned task, which learns a new but related task efficiently.

To verify the last advantage of memory property, after all the RL decoders achieved a good performance on the second task, we trained the decoders back to the first task. The learning curves are shown in Fig. 4d. When reverting to the first task, the baseline RL decoder with a neural network structure (blue) and the KRL with no alignment decoder (green) undergo a notable performance drop, despite the prior training on the first task. Conversely, CATKRL based decoder (red) maintains good performance with almost no performance drop, which demonstrates the decoder’s retention of task 1 following the training of task 2. The reason is shown in Fig. 4e with the neural data distribution and the corresponding weights by different decoders. Once the decoders are well-trained in the two-lever task, both CATKRL and clustering KRL have stored the neural patterns and the weights to the corresponding actions. When reverting to the one-lever task, the decoders would choose actions based on their fresh parameters learned from the two-lever task. Without neural pattern alignment (right figure, Fig. 4d), the neural patterns between two tasks are mismatched. The action selection could be different from the ground truth action labels (red hollow bar on green dot clouds), which decreases the decoding performance. As our method has aligned the neural patterns in the most similar cluster between two tasks, the decoder could highly likely choose the correct action (solid bar centered at neural data cloud with the same color) utilizing the shared neural-action mapping across two tasks, which maintains the decoding performance of CATKRL when switching back to the one-lever task. A similar phenomenon is also observed in the testing performance over multiple subjects (t-test, p<0.05) as shown in Fig. 5b.

The memory property of the CATKRL decoder can also be viewed from data efficiency (Fig. 4f). All RL decoders were first trained from the one-lever task to the two-lever task as the initial condition. Then the decoding performance was tested with different amounts of training samples when reverting to the one-lever task (Fig. 4f). Compared with the baseline RL decoder (blue) and the KRL decoder with no alignment (green), we observe that CATKRL (red) has a significantly higher testing success rate when there are small amounts of training samples (≤40%). Even though the final performance values of the decoders are similar, CATKRL requires fewer training samples (40%) to achieve optimal performance. CATKRL applies the clustering alignment in neural patterns and requires the least amount of data (40%) to reach the best decoding performance. These results suggest that CATKRL could efficiently retain the performance of the old task even after the training of a new task.

In summary, we posit the existence of a transfer mechanism in the brain that facilitates motor learning. This mechanism recognizes similar neural patterns and aligns the corresponding neural-muscular mapping for new task learning. We visualized the process of neural pattern development within the feature space by an iterative linear alignment. And we computationally echoed the neural pattern evolution by alignment iteration. Furthermore, the transfer principle is incorporated into the design of a kernel reinforcement learning based decoder. The evaluations on the decoding performance demonstrate that our algorithm could utilize the hypothetic brain transfer mechanism to enhance the learning ability of BMI. This is shown by the faster learning speed with less training data in the new task, also a retained performance in the learned task.

## Discussion

In this study, we investigated a potential motor learning transfer mechanism at the neural ensemble level: during the learning of a new task, the neural patterns of the same action tend to cluster, and the shape of the cluster and the centroid distance remain similar across tasks. The neural patterns could be aligned in feature space. Finally, the established neural-action mapping is transferred through the aligned neural patterns to achieve more efficient learning in the new task. Based on this hypothetic mechanism, we proposed a transfer learning algorithm (CATKRL) that utilizes such mechanism to facilitate the learning ability of BMI on the new task. Our experiments demonstrated that the subjects transferred the learned movements from the simple task to the related complex task, which is shown by the decreasing response time and increasing success rate (Fig. 1c and 1d). Our study also revealed the neural pattern development of motor learning transfer in the feature space using jPCA (Fig. 3a).

We note that, in support of the transfer mechanism, our results only present the alignment in the feature space instead of the original neural firing activity space. Our simulation demonstrates that the neural pattern evolvement in the linearly projected low-dimensional space could reflect the neural population dynamics of the original spike activities. If the alignment in the feature space is noticeable, it can be inferred that the alignment also happens in the neural ensemble space.

Previous studies have shown that low-dimensional latent neural activities remain stable during a long time of training for the same task ^13,28^. We assume that a similar scenario happens in the feature space visualized by jPCA, and the stability is reflected in the maintenance of the shape and the centroid distance of the neural pattern clusters corresponding to the learned movement (Fig. 3a). The neural pattern alignment could transfer the acquired neural-muscular mapping to facilitate the learning of the new task.

Another contribution of our study, in terms of computational tools, is the proposal of a novel algorithm (CATKRL) that tries to utilize the transfer mechanism from the brain, trying to enhance the learning ability of BMI. First, we cluster the neural patterns in the feature space to build the reservoir that relates to all kinds of action (Fig. 6b). Through the training of CATKRL by trial and error, each neural cluster gradually establishes the weights to corresponding actions, which represents the learned neural-muscular mapping. Second, CATKRL evaluates the similarity of pair-wise clusters between two tasks, which allows the identification of the most similar clusters. Finally, CATKRL aligns the most similar cluster pair and transfers the weights from the old cluster to the new one with the most similar shape. The decoding in the new task is supported by the transferred parameters within the similar cluster, while other parameters in the existing clusters are reserved. In this case, CATKRL could help the learning of the new task with a fast-learning curve and less training data.

**Fig. 6.**
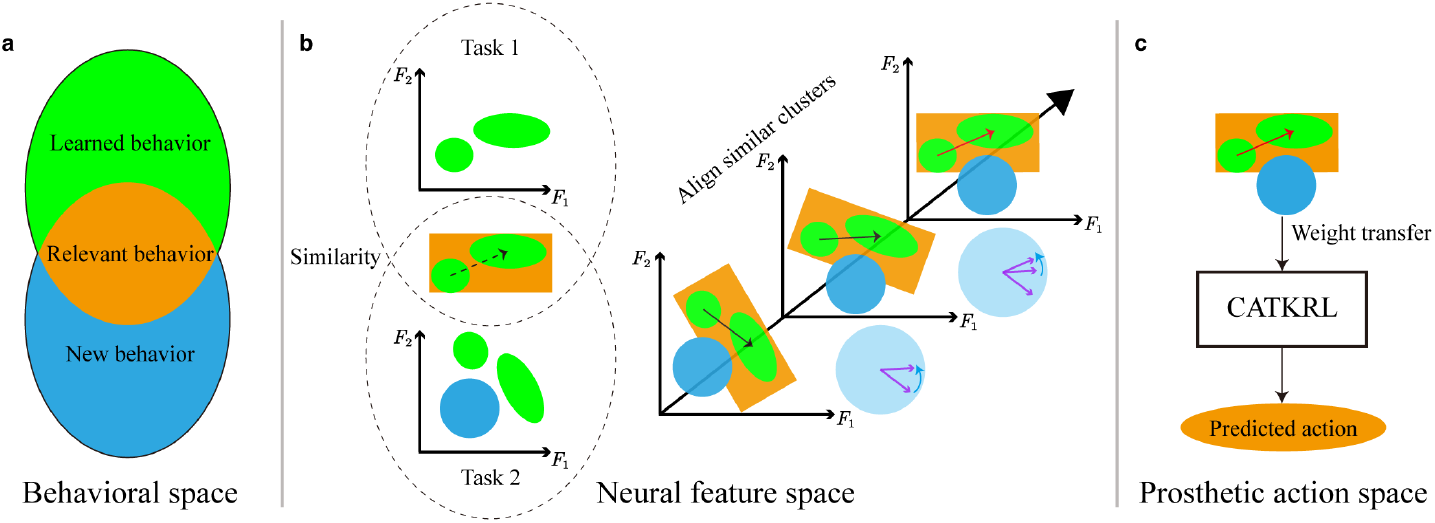
The clustering alignment method detects and exploits shared neural-action mapping from neural pattern clusters between two tasks. **(a)** The association between two tasks in the behavioral level. During the BMI training, the animals were first instructed to learn a relatively easy task. The behavior in task 1 is shown as the green ellipse. After the animals were well trained, they were guided to learn a new task which is harder but related to task 1. The required behavior of task 2 is shown in the blue ellipse. The two tasks are designed to have some relevant behaviors (orange) so that the animals can re-utilize the actions learned from task 1. **(b)** The observation of the animal neural activities in the feature space. We project the high-dimensional neural activities into a low-dimensional feature space. The neural patterns tend to form clusters for the same actions, shown by the green circles. Since there are new behaviors in task 2, there is one more cluster that appears in the new task, which is shown by the blue circle. The positions of the neural patterns for the learned behavior in task 2 have changed. But the shapes of the green clusters remain similar in task 2, which gives us a way to identify the neural patterns corresponding to the relevant behavior. Based on each cluster’s covariance matrix, we could measure the pairwise similarity between clusters in two tasks. And the most similar clusters are labeled within the orange rectangle. Then the decoder would rotate and shift the entire neural patterns in task 2 (blue and green clusters) to iteratively align the coordinate of clusters (green) in task 2 in order to match (the black vector) as close as to the most similar clusters (green) in task 1. The light blue plate shows the rotational trajectory of all iterations. The purple arrow represents the direction of the relative position between two green clusters. The blue arrow represents the counterclockwise rotation. **(c)** The aligned neural patterns are input into a clustering-based kernel reinforcement learning to probabilistically choose the corresponding actions. Since the decoding method has been trained in task 1, it has already stored the weights that map the neural patterns to the corresponding action in green clusters. The decoder could utilize the weights from task 1 to translate the neural patterns in task 2 into the corresponding action efficiently. Based on the reward signal through trial and error, the weight of CATKRL is updated. The coordinate of the new pattern in task 2 is also stored in the algorithm to follow the new patterns in task 2.

One important thing to notice is that in our experiment, even though the tasks are different, part of the actions are consistent across tasks, which means we only show one particular instance of motor learning transfer. Motor learning transfer might also be achieved by a combination of the learned actions. Thus, the generalization of our results for the motor learning transfer is still an open question. For example, a complex behavior could be decomposed into a set of simple behaviors. We presume that the neural pattern in the brain could also be decomposed into small clusters and combined in a new sequence for the new task to achieve motor learning transfer. CATKRL could be a promising tool to deal with this scenario in BMI. During the learning of a new task, CATKRL could identify and associate multiple clusters that are similar to the current task. Then CATKRL could transfer the parameters of the neural action mapping from the identified clusters to the new task. The association and transfer procedures are iterated during the algorithm’s online training through trial and error. Hopefully, CATKRL could enrich the learning ability of BMI in this complex task. In the future, we could design the experimental paradigm so that the new task requires a combination of the actions in the learned task. In this case, we could apply CATKRL to study the more complex transfer mechanism in the brain, as well as build a more general decoder to improve the learning ability of BMI.

Our proposed novel algorithm CATKRL could be a valuable tool for clinical applications in BMI. For example, if a paralyzed patient wants to restore arm function, he/she could translate the brain signal to control the prosthetic arm by BMI. The ultimate goal is to let the patient control the prosthetic arm dexterously as his/her real arm, which contains all kinds of movements during the daily interaction and much beyond the experiment conducted in the lab. To achieve the ultimate goal, we could decompose complex movements into a set of simple tasks (e.g. move the arm up/down/left/right). The patient could start with the simple task to accumulate experience on the prosthetic control. At the same time, through trial and error, CATKRL builds up the reservoir of the neural pattern clusters and accumulates the neural-action mappings from simple movement to complex tasks. As the patient starts the adaptation from the simple task training (e.g. directional moving) to a more complex task (e.g. draw a circle). His/her brain could search for similar experience from the simple task to assist the learning in the more complex task. Meanwhile, CATKRL-based decoder would take the neural inputs during the subject’s search and exploration, align and transfer the neural-action mapping from similar neural clusters to better interpret the patient’s movement intention. In this way, when we design the BMI task training in the sequence of incremental difficulty, through the co-adaptation between the patients and the CATKRL decoder, the patient might achieve the full restoration of motor control on the prosthetic arm as his/her real arm.

In conclusion, our proposed algorithm CATKRL could advance the understanding of the brain’s transfer mechanism during motor learning, especially how the brain encodes and utilizes the learned action. On the other hand, CATKRL also provides a promising tool that could enrich the learning ability of BMI, which includes a faster learning speed with less training data on the new task, and the performance maintenance on the old task. It opens the possibility to enable BMI for the full motor restoration for clinical trials.

### Experiment and Subject Details

The animal training and electrode implantation of the male Sprague Dawley (SD) rats were conducted in the Hong Kong University of Science and Technology (HKUST). All the procedures were approved by the Animal Ethics Committee at HKUST. The SD rats were implanted with two 16-channel microelectrodes (15 mm electrodes, spaced 200 µm apart). One in the primary motor cortex (M1), another in the medial prefrontal cortex (mPFC) of the left hemisphere, which was contralateral to the lever-pressing forearm in the task. The rats recovered at least one week prior to these experiments. During the movement task, the neural signals were recorded by Plexon (Plexon Inc, Dallas, Texas) with a 40 kHz sampling frequency. Then the voltage signals were passed through a 4-pole Butterworth high pass filter (500 Hz). Then the signals were processed to detect spikes by thresholding at -4σ where σ is the standard deviation of the signal amplitude. The neural data were sorted by the automatic valley seeking method ^29^. The cue and movement events were recorded using a behavior system from Lafayette Instrument, USA. The event signals were synchronized with the neural signals.

In this experiment, the rats performed a one-lever pressing task for the first several days until well-trained, then started to learn a two-lever discrimination task as described in Fig. 1a and 1b. At the beginning of each trial in the one-lever task, an audio cue (10 kHz, duration 0.9 s) was presented to the rat. The rat needed to press the lever in the behavior box within 8 seconds. Otherwise, this trial was considered a failure. When the rat pressed on the lever, it needed to hold it for 0.5 s until a success audio cue (10 kHz, duration 0.09 s) was given. Then a drop of water would be given to the rat as the reward for this successful trial. The inter-trial interval is randomly chosen within 4∼6 s. The rats were trained on the one-lever pressing task for two weeks on average. The response time and success rate show that the rat gradually became proficient in the task (Fig. 1c and 1d). After the rats were proficient in the one-lever pressing task, they would learn to perform a more complex two-lever discrimination task as shown in Fig. 1b. The configuration of the behavior box was almost the same as the first task. But there was one more lever (low lever) below the first lever (high lever).

When the trial started, there would be two different audio cues (0.9 s), which were 10 kHz (high cue) and 1.5 kHz (low cue) respectively. The rats needed to distinguish the audio cues and press the corresponding lever. If it was a high cue, to get the water reward, the rat must press and hold the high lever for 0.5s, which was the same as the first task. Thus, the rats might utilize the experience of pressing the high lever in the new task. If the trial started with a low cue, the rat should press and hold the low lever (orange) for 0.5 s. Then it could get a drop of water reward. When the rat did not press any lever within 8 seconds or pressed the wrong lever, it could not get the reward. The inter-trial interval was randomly chosen within 4∼6 s. The response time and success r”te s’ow that the rat gradually became proficient in the two-lever discrimination task (Fig. 1c and 1d).

## Methods

In this study, we propose a novel Clustering Alignment-based Transfer algorithm on Kernel Reinforcement Learning (CATKRL) to model the neural pattern evolution during task learning, which utilizes a possible transfer mechanism on the neural level during motor learning to enable efficient task learning. The structure of the proposed algorithm is shown in Fig. 6. We first project the high-dimensional neural activity into a low-dimensional feature space (e.g. jPCA, Fig. 6b), where the neural patterns are observed to group together for the same action. During the training of the first task, our proposed decoder will transfer the learned neural-action mapping to more efficiently control prosthetic movements in a new movement task. The transfer procedure to the new task is shown as follows. First, our proposed algorithm explores the neural patterns among two tasks and identifies the neural clusters with a similar shape. During the learning procedure of the new task, we observe some clusters in neural patterns have a drift in the feature space, but the cluster corresponding to the same action maintains a similar shape (green clusters). Second, we iteratively align the neural clusters from the new task to the old task. The alignment iteration illustrates a dynamical process of how the similar neural pattern is identified, which is provides a possibility to explain how the brain finds learned experience. Finally, the aligned neural patterns are input into a kernel reinforcement learning scheme to control prosthesis to complete the new task (Fig. 6c). To re-utilize learned neural-action mapping, CATKRL transfers the decoder parameters in the old task when interpreting neural-action mapping in the new task, which increases the learning efficiency. Taking together all the steps, CATKRL reveals and exploits the potential transfer mechanism of the brain to enhance task learning in BMI. The details of the proposed algorithm would be shown as follows.

## Projection to Low-dimensional Feature Space that Captures Rotational Structure

The populational neural activity is represented by a high-dimensional vector ***u***_*t*_ ∈ *R*^*P*×1^ , where each component represents the spike count within the time window *t* (the length of the window is 100 ms). *P* is the dimension of the neural input, which contains the total number of recorded neurons (*N*) and the number of historical embedding ( *H* ) of the neural activity (*P* = *N* + *N* ⋅ *H*). For the rat data, *N* = 32, *H* = 5, *P* = 192.

The vector of neural activity is high-dimensional, which makes it challenging to interpret the neural population dynamics. We first reduce the dimensionality of the neural activities to *P*′ by PCA to reveal the most significant variation in the neural data (*P*^′^ = 5). Then we apply a method called jPCA ^26^ to best capture the rotational structure in the data, which works as follows. First, the neural pattern after PCA is modeled by the following equation 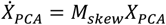, where *X*_*P*CA_ is the data matrix after PCA and 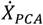 is the derivative of the data matrix. *M*_*s*kew_ is a skew-symmetric matrix that describes the relationship between the data matrix and its derivative. *M*_*s*kew_ has only imaginary eigenvalues and the most significant rotational dynamics happen in the plane defined by the eigenvectors with the largest complex-conjugate eigenvalue pairs, denoted by *V*_1_ and *V*_2_ . Since the eigenvectors are imaginary, the real jPCA axes are calculated by jPC_1_ = *V*_1_ + *V*_2_ and jPC_2_ = j(*V*_1_ − *V*_2_). The rest dimensions of the jPCA space are calculated by the similar eigenvector operation. Finally, the neural data after PCA are linearly projected into jPCA space (*D* = 3) to illustrate the rotational dynamics.

## Identification and Alignment for Similar Neural Pattern Clusters

In the jPCA space, we propose to apply the Density-Based Spatial Clustering of Applications with Noise (DBSCAN) ^30^ as the clustering method to better reflect the neural pattern distribution. The first advantage of DBSCAN is that it groups the neural patterns in the dense region, which forms a compact representation for the neural patterns of the same action. Another advantage is that DBSCAN would mark the low-density region as outliers, which ignores some of the neural patterns that might correspond to the noisy movement. After applying DBSCAN on the neural patterns, the *i*^th^ cluster in the first and second task are denoted as 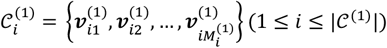 and respectively. 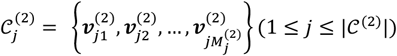 and |𝒞^(2)^| represent the total number of clusters of two tasks respectively 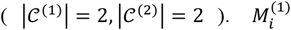 and 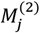 represent the total number of neural data points within cluster *i* and *j* in the first and second task respectively.

The neural pattern cluster could be described by the centroid and the expansion across dimensions. We represent each neural cluster as an ellipse centered at the mean value of the neural patterns. And the principal axes of the ellipse show the directions where the neural pattern varies the most. We run the Principal Component Analysis (PCA) for each cluster 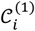 and 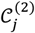. The eigenmatrices of the first and second task are denoted as 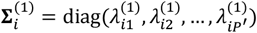 and 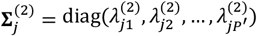 respectively, where the diagonal elements are the eigenvalues sorted in descending order. Each eigenvalue shows the degree of extension along the corresponding eigenvector. We then compare the pairwise Covariance Matrix Distance ^31^ between the clusters of the first task and the second task, which is shown in the following equation

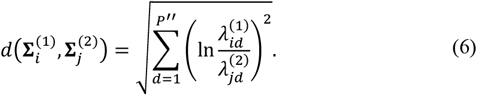

By comparing the pair-wise Covariance Matrix Distance between clusters in two tasks, we could find the cluster pair with the smallest Covariance Matrix Distance, denoted as 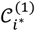 and 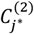. These two clusters have the most similar shape, which presumably contain neural patterns corresponding to the similar action (red cluster in Fig. 3a). We could use 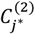 and 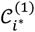 as the alignment source and target respectively.

The next step is to match the neural patterns between two clusters as close as possible to re-utilize the learned neural-action mapping in the first task. We first make the neural patterns to be centered at the origin to do the rotation. The zero-mean clusters are denoted as 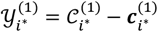 and 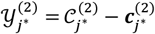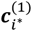, where 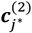 and ***c***^(2)^ denote the centroid for cluster *i*^∗^ and *jj*^∗^ respectively. For every neural data point in feature space 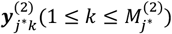 in 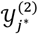, based on the distance to all the neural patterns in 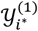, we can find its closest neural pattern 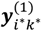 in 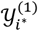. The closest neural patterns will form a set denoted as 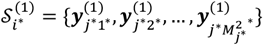. One thing to note is that different elements in 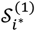 might refer to the same neural pattern in 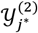 . Now the specific goal becomes matching the two neural pattern sets 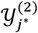 and 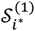 as close as possible, which could be accomplished by the one-step rotation matrix **Ω**^(*n*)^ from solving the Procrustes problem ^32^ as follows

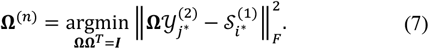

Here 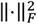 denotes the square of the Frobenius norm, which is the summation of the squares of all the elements within the matrix. The goal of the rotation is to minimize the overall distance of the closest neural patterns between 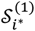 and 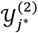. The Procrustes problem could be solved in a closed-form ^32^. After getting the rotation matrix **Ω**^(*n*)^ , we need to re-associate the nearest neural patterns between 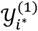 and 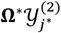.

Then we will update the element in 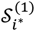 by the newly associated nearest neural patterns. And a new rotation matrix **Ω**^(*n*+1)^ would be calculated from (2). This iteration continues until there are no more changes in 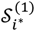. In each iteration, the source neural pattern cluster would get closer to the target cluster as shown in Fig. 3b.

We get the final **Ω**^∗^ to align the two most similar clusters until the iteration stops. Since the relative position of the clusters within one task remains similar (black arrow in Fig. 3a and 3b), we could use the same matrix to align all neural patterns between the two tasks. We denote 𝒰^(1)^ and 𝒰^(2)^ as the sets that contain neural patterns in the old task and the new task respectively. ***c***^(1)^ and ***c***^(2)^ are the centroids of 𝒰^(1)^ and 𝒰^(2)^ and respectively. Finally, the neural patterns in the second task are aligned to the first task by the following equation

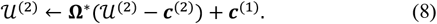

## Efficient Decoding in the New Task by Weight Transfer

We proposed a Clustering Alignment based Transfer on Kernel Reinforcement Learning (CATKRL) to utilize the transferred neural-action mapping on the aligned neural patterns. The first advantage of our method is that it inherits the property of reaching the global optimum in the kernel reinforcement learning methods, such as QAGKRL ^33^ and CKRL ^34^ , which ensures a smooth transfer in the more complex task. The second advantage is that our method directly operates on the neural cluster after alignment and decodes the neural patterns in the subspace with efficiency and also provides the ever-growing feature space to store the neural patterns explored in task learning. The input of CATKRL is the neural patterns in the feature space after alignment and the output is the corresponding action of the subject. CATKRL is trained by the neural-action data from the simple task to the more complex task.

When the training for the second task starts, the decoder needs to build a neural-action mapping based on the first task. At the time window *t*, the current neural pattern after alignment is denoted as 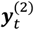 . The corresponding neural-action mapping is written as

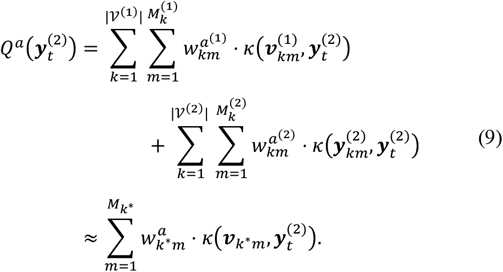

The corresponding action value 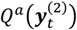 consists of two components. The left part is the neural-action mapping learned from the first task. After the training of the first task, CATKRL clusters the neural patterns as 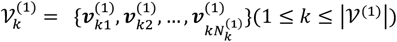 (Fig. 6b, first row, circles). |*V*^(1)^| is the total number of clusters and 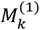 is the total number of neural patterns within cluster *k* in the first task. 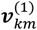 and 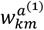 are the *m*^th^ neural pattern from the *k*^th^ cluster in the first task and its weight connected to the *a*^th^ action respectively. 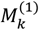 is the total number of neural patterns within the *k*^th^ cluster in task 1. The right part in Equation (9) represents the neural-action mapping that is newly learned in the second task. The symbols have similar meanings as the left part except that the neural patterns 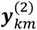 and weights 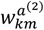 newly appeared in the second task.

The action value is calculated by the weighted sum of the Gaussian kernels across clusters in two tasks, where 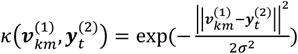 . *σ* is the kernel size. Due to the exponentially decayed tail of the Gaussian kernel, we can effectively use the nearest cluster to do the decoding as shown by the second line in (9). *k*^∗^ is the nearest cluster to the current neural pattern 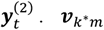 and 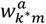 are the 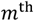 neural pattern from cluster *k*^∗^ and its weight connected to the *a*^th^ action respectively. 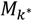 is the total number of neural patterns within cluster *k*^∗^ . For the current neural pattern 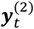 after alignment, if the nearest cluster is already formed the first task, decoder could re-utilize the learned weights, which increases the learning efficiency. On the other hand, if the nearest cluster newly appears in the second task, the decoder is able to explore the new neural-action mapping.

After the action values are calculated, the action is decoded probabilistically based on the softmax policy. If the chosen action is the same as the subject’s true action, the neural-action mapping will be reinforced by setting the reward signal *r* = 1; otherwise *r* = 0 and the neural-action mapping is punished. The parameters are updated through trial and error. More details of the action selection and weights update could be found in ^34^.

## Acknowledgements

This work was supported in part by the Science Technology Innovation (STI) 2030-Major Projects under Grant 2021ZD0200403; in part by the Research Grants Council of Hong Kong Special Administrative Region, China, under Project Hong Kong University of Science and Technology (HKUST) C6049-24G; in part by the Special Research Support from the Chau Hoi Shuen Foundation under Grant R9051; in part by the Innovation and Technology Commission under Grant ITCPD/17-9; and in part by the Seed Fund of the Big Data for Bio Intelligence Laboratory from HKUST under Grant Z0428.

